# Molecular characterization of spiny lobsters from the Saint Martin’s Island, Bangladesh: a new record of *Panulirus longipes* (A. Milne-Edwards, 1868)

**DOI:** 10.64898/2026.05.29.728916

**Authors:** Sadia Afrin Kamal, Nur-A-Zannat, Umme Khadiza, Kazi Ahsan Habib

## Abstract

In Bangladesh, spiny lobsters are vital to the economy as a key export. A total of 4 species of spiny lobsters from the genus Panulirus have been identified in Bangladeshi waters (*Panulirus homarus, P. ornatus, P. polyphagus and P. versicolor*). This study aimed to identify and update the existing list of spiny lobster species found in Bangladeshi waters by examining their morphological characteristics and the phylogenetic profile of the cytochrome c oxidase I (COI) from mitochondrial DNA (mtDNA) gene marker. A new species, *Panulirus longipes*, was recorded for the first time from the Saint Martin’s Island, Bay of Bengal, Bangladesh. This species is characterized by white spots and longitudinal orange stripes on its walking legs. In the Neighbor-joining (NJ) tree, the sequences of the same species were grouped together under a single clade for COI and demonstrating the effectiveness of marker genes in distinguishing between lobster species. The results indicate a new lobster species in Bangladesh, enhancing the known diversity of lobsters in the region and revealing previously undocumented species.

## Introduction

Crustaceans are a large group of arthropods, 4^th^ largest diversity among the animals group and are usually considered as a subphylum (Holthuis, 1991). There are between 50,000 and 75,000 species in this category, including many well-known creatures like crabs, lobster, barnacles, prawns, woodlice, and beach fleas, in addition to several lesser-known species (Anger, 2001). Lobsters, highly valued crustaceans, can be found in tropical, subtropical and temperate water (Radhakrishnan et al., 2019). Approximately 149 lobster species have been identified in the world (Holthuis, 1991). The two families that comprise the majority of lobster species are the the Scyllaridae (slipper lobsters) and Palinuridae (spiny lobsters) (Ahmed et al., 2022). Around the world, lobster from the Palinuridae family is regarded as a delicious seafood variety (Senevirathna and Munasinghe, 2013). The Palinuridae family consists of 47 species of the genus palinurus while the Scyllaridae family comprises 88 species, 19 genera, and two subspecies worldwide (Holthuis, 1991; Carpenter and Niem, 1998; Chan, 2010; Yang et al., 2011; Yang and Chan, 2012; Ahmed et al., 2022). In Bangladesh, 4 species of spiny lobsters (family Palinuridae) and 2 species of slipper lobsters (family Scyllaridae) have been reported (Hossain, 2001; Ahmed *et al*., 2022; Habib *et al*., 2023).

The long-legged spiny lobster, *Panulirus longipes*, is a member of the Palinuridae family and is distinguished by the large number of round white spots covering its body, particularly the abdomen (Liu and Pu, 2021). This species is indigenous to the tropical and subtropical Indo-Pacific region, and its range includes Madagascar, the east coast of Africa, the Mediterranean, India, Malaysia, Japan, Taiwan, the Philippines, Indonesia, Papua New Guinea, and northern Australia (George and Rao 1965; Pillai and Thirumilu 2007; Spanier and Friedmann 2019). They are usually found in clear or slightly turbid water at depths between 70 and 400 m (George, 2005; Liu and Pu, 2021). In addition to consuming mollusks and other marine invertebrates that inhabit the seabed, they are nocturnal and tend to be reclusive. They are caught more often due to their high nutritional value, strong market demand, and competitive prices; they are sold both fresh and frozen in local markets or as major exports (Liu and Pu, 2021).

In morphological analysis, the exoskeleton is a crucial characteristic for identifying lobsters (Akmal et al., 2023; Irfannur et al., 2024). But Phenotypic morphological characterization can sometimes result in unsuccessful or inconclusive identification of crustacean species due to variations in body color and polymorphisms influenced by diet, growth, habitat, seasons, and morphological similarities among sympatric species, as well as the effects of rough handling (Rajkumar et al., 2015; Ahmed et al., 2021; Kaur et al., 2021; Yusnaini, Muzuni & Nur, 2021). Furthermore, the genus *Panulirus* represents a valuable global trade commodity, with 21 recognized taxa (Ptacek et al., 2001), all of which are considered threatened (IUCN, 2023) due to excessive overfishing and inadequate conservation efforts. Regular monitoring of *Panulirus* species is crucial to developing effective management strategies. Additionally, accurate species-level identification is essential for quality control in the food industry to ensure proper labeling of traded crustacean products (Kappel et al., 2021) and to distinguish processed seafood products (Ahmed et al., 2022). There is also limited information on the species availability and genetic diversity in Bangladesh (Ahmed et al., 2021). In this context, molecular techniques are vital for accurate species identification and understanding evolutionary relationships among various marine taxa (Radhakrishnan, Kizhakudan & Phillips, 2019). Molecular data are especially beneficial when morphological and genetic information is inconsistent (Palumbi et al., 1991; Sarver, Silberman & Walsh, 1998). Combining morphological and molecular approaches can improve the efficiency of biodiversity assessments (Jamaluddin et al., 2019; Alam et al., 2021). Therefore, given the limitations of morphology-based identification, there is a need for new taxonomic recognition methods, such as DNA-based identification techniques (Hebert et al., 2003).

DNA barcoding is a novel approach that uses short, standardized gene sections of the mitochondrial Cytochrome Oxidase subunit I (COI) gene to offer quick, precise, and automatable species identifications (Herbert & Gregory, 2005; Casiraghi et al., 2010). The process of DNA barcoding facilitates the quick identification of new species, enhances the accuracy of taxonomic data, and makes data easily accessible to non-taxonomists and researchers in general (Hebert et al., 2003; Stoeckle, 2003). Barcode sequences could be made more readily available for species identification and biodiversity study by being deposited in a public database along with primer sequences, trace files, and quality scores (Lorenz et al., 2005).

Limited studies have been conducted in Bangladesh on the molecular identification and comprehensive taxonomy of spiny lobsters. Under the family Palinuridae, only four species of spiny lobster (*Panulirus homarus, P. ornatus, P. polyphagus and P. versicolor*) have been identified thus far from Bangladesh (Ahmed et al., 2022). However, the aim of this study is to identify *P. longipes* for the first time from Saint Martin’s Island, Bangladesh, based on morphomeristics as well as mitochondrial cytochrome c oxidase subunit I (COI).

## Materials and Methods

### Sample collection and preservation

Lobster specimens were collected from Saint Martin’s Island (20º 07.225’N, 92º 11.420’E) in Bangladesh between December 2023 and February 2024. The island is situated at the southern tip in the Bay of Bengal, Bangladesh (Fig). After tagging, the samples were photographed using the method outlined by Randall (1961). The samples were then sent to the Aquatic Bioresource Research Lab (ABR Lab) at the Department of Fisheries Biology and Genetics, Sher-e-Bangla Agricultural University, Dhaka, Bangladesh, for analysis. Following morphological examination, a small piece of muscle tissue was taken from each specimen and preserved in a sterile 1.5 ml tube with 99% alcohol for future molecular analysis. All specimens examined were stored in the ABR Lab for additional research.

### Morphological Observations

Morphological study was conducted of the lobsters to identify these species, using methods that are described by Weber and de Beaufort (1916), Mabuchi et al. (2014), and Psomadakis et al. (2019).

### Genomic DNA extraction, PCR amplification and, sequencing

Genomic DNA was isolated from the collected tissue samples using TIANamp Marine Animals DNA Kit (TIANGEN) following the instructions provided with the kit. The concentration of the extracted genomic DNA was quantified using a Qbit 3.0 fluorometer. Polymerase Chain Reaction (PCR) was performed in small reaction tubes (0.2 ml) using a Thermal cycler (2720 Thermal Cycler, Applied Biosystems) with a total of 25µl reaction mixtures. The COI gene fragment of the mitochondrial DNA (mtDNA) was amplified using the universal primer set for crustaceans LCO1490 (forward) 5’-TCAACAAATCATAAGGACATTGG-3’ and HCO2198 (backward) 5’-TAAACTTCAGGGTGTCCAAAGAATCA-3’ (Folmer *et al*. 1994). The PCR thermal cycling conditions consisted of pre-heating at 95ºC for 5 min, followed by 40 cycles of denaturation at 94ºC for 45s for, annealing at 42ºC for 30s, extension at 72°C for 45s and completing with a final extension at 72ºC for 10 min. Following the successful PCR amplification, each sample was visualized on 1% agarose gel (EZ-Vision® In-Gel Solution, USA) and stained with ethidium bromide. The gel documentation chamber (Model: Syngene InGenius3) equipped with UV-ray light was used to observe the DNA bands on a connected computer using GeneSys Software. PCR samples showing clear, single bands were subjected to purification using the PCR Purification Kit (TIANGEN-Universal DNA Purification Kit) for subsequent sequencing. The concentration of the purified DNA was determined by Qbit 3.0 fluorometer. Purified samples were sent to Macrogen Inc. (Korea) where sequencing was carried out using the Sanger method with an automated sequencer (ABI 3730 × 1 DNA analyzer).

### Molecular analysis

The DNA sequences obtained were edited for clarity based on coherent peaks in the chromatogram, using Chromas Lite 2.1 and Geneious 9.0.5 (Kearse et al. 2012), along with manual verification. Start and stop codons in the COI sequences were examined using the Expasy translation tool (Artimo et al. 2012). Each sequence was validated through a BLASTn search against the nucleotide database, focusing on the best matching sequences with an identity threshold of over 99%. All sequences were aligned using ClustalW (Thompson et al., 994). The nucleotide composition of the COI sequences for each species was analyzed using MEGA X (Kumar et al. 2018). Pairwise genetic distances and the overall mean distance were calculated using the Kimura-2-parameter (K2P) model (Kimura, 1980) in MEGA X. The phylogenetic position of the species studied within the family Palinuridae was assessed using the Neighbor-joining (NJ) method in MEGA 11. The analysis included 25 additional species from the same genera representing COI sequences from various regions, along with an out-group from the family Scylaridae (Thenus genus). The robustness of the phylogenetic relationships was tested through bootstrap analysis with 1,0000 replications.

## Results

### 1. Morphological analysis

***Panulirus longipes*** (A. Milne-Edwards, 1868)

#### English Name

Long-legged spiny lobster

#### Material examined

Specimen collected from St. Martins Island, Bay of Bengal, Bangladesh (20º 07.225’N, 92º 11.420’E); February 2024; 1 Male; Total Length (TL): 49.5 mm; Carapace Length (CL): 9 mm; Carapace Width (CW): 4.5 mm; Body Weight (BW): 377g.

#### Habitat

Inhabits clear or slightly murky waters at depths ranging from 1 to 18 meters. Typically, in rocky environments and coral reefs.

#### Diagnostic characteristics

Both carapace and abdominal regions are covered with disorderly dispersed white and orange spots. The body and the post-orbital area are brown, while the dorsal parts of the carapace and anterior segments of the abdominal region are dark brown in color. The base of the antennal peduncle is violet, and its inner side of dorsal region is pinkish brown. The antennular plate’s frontal margin is armed with two separate small spines. The carapace carries numerous spines of different sizes; some are black, while some have orange tips with white bases. Antennular flagella are cross-banded in white. Legs bear orange and white spots at the end of each segment. Transverse grooves are present in every abdominal segment which is connected laterally to the pleural grooves. Exopods are appeared in the second and third maxillipeds. Tips of legs bear reddish brown hairs (Fig 2). The comparable morphological characteristics that are utilized to identify spiny lobsters are displayed in Table 2.

**Fig 1:**
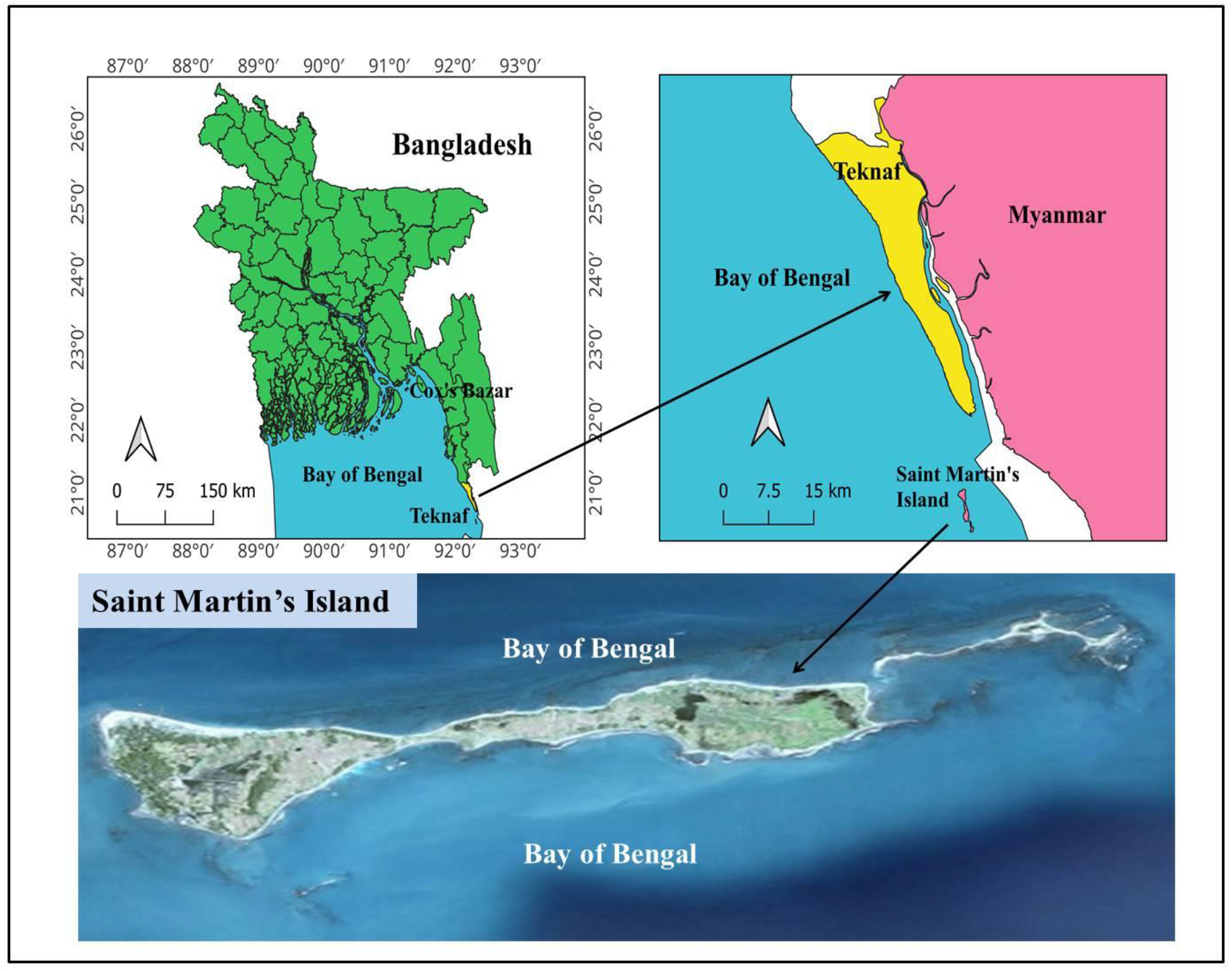
Location of the Saint Martin’s Island of Bangladesh in the northern Bay of Bengal from where fish samples were collected.

**Fig 2:**
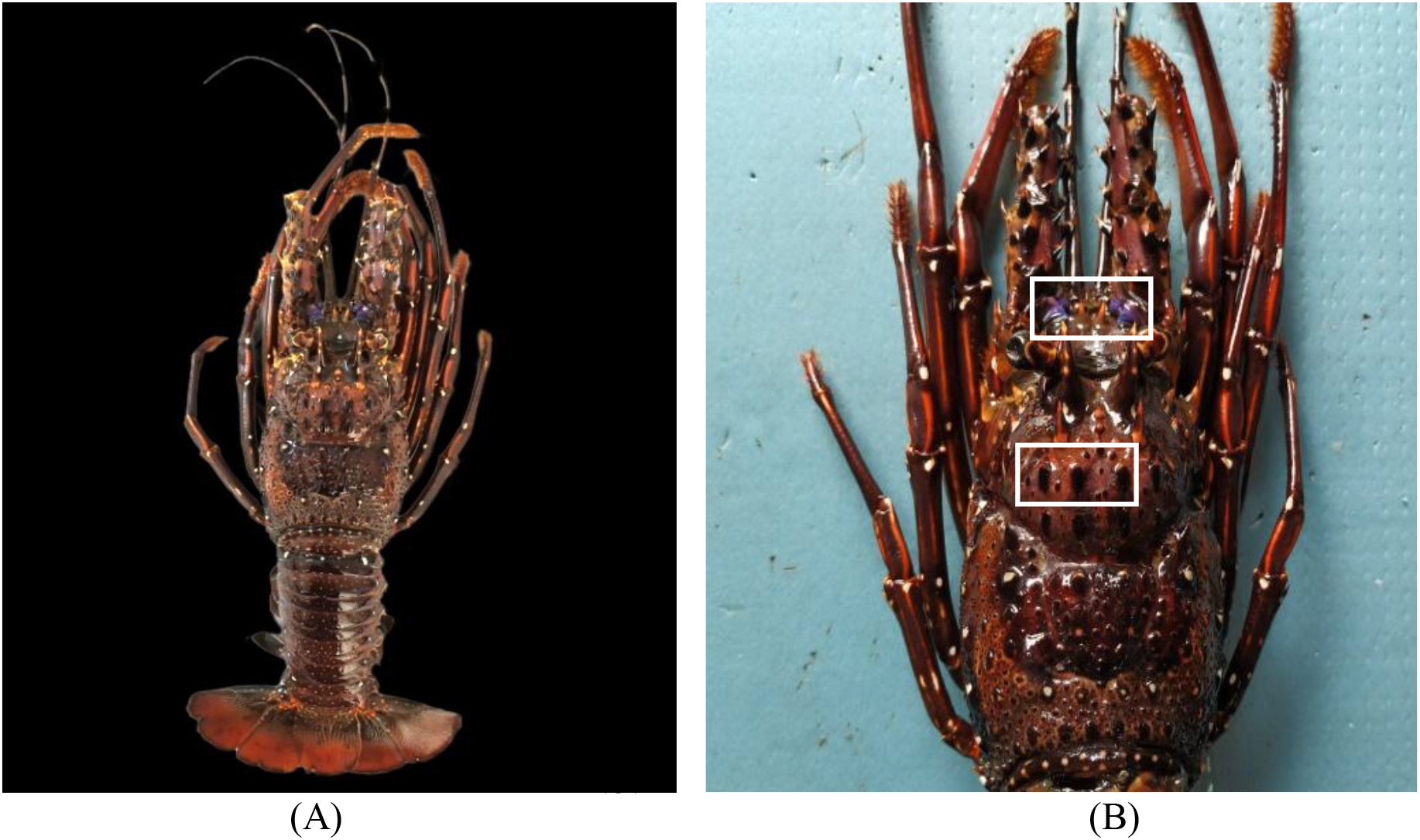
**(A)** The longlegged spiny lobster (*Panulirus longipes*, A. Milne-Edwards, 1868) collected from Saint Martin’s Island, Bangladesh, **(B)** Identification of *P. longipes* based on the presence of 3 spines in a row and purple antennular peducle.

**Fig 3:**
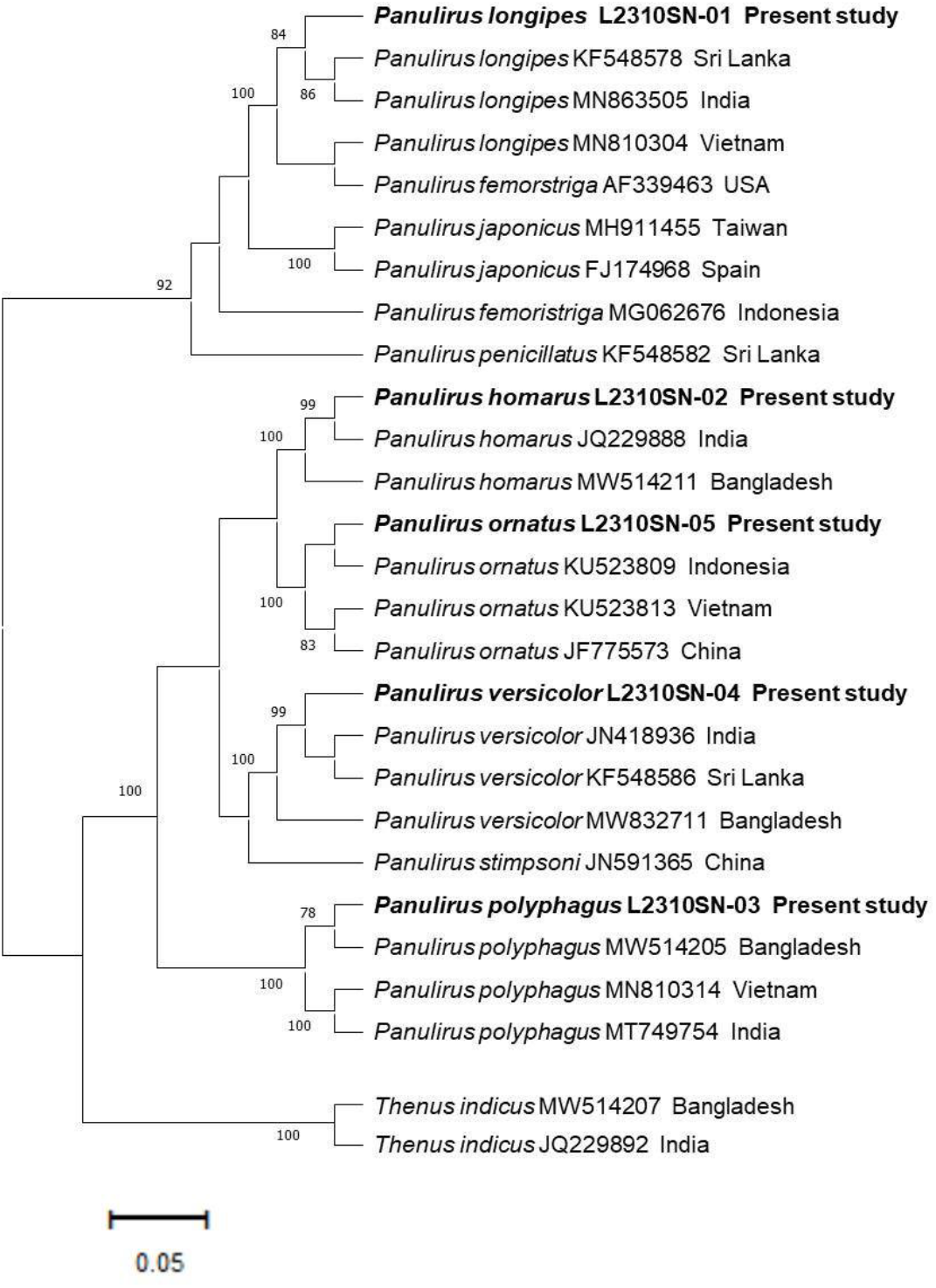
Neighbor-joining (NJ) tree inferred from the partial mitochondrial COI sequences of 25 *Panulirus* species, with *Thenus indicus* as out-group. Bootstrap values of >70% are shown.

### 2. Molecular analysis

Five COI barcode sequences of five different Panulirus species were generated, with base pairs ranging from 634 to 652. A total of 27 species from the families Palinuridae and Scyllaridae, including *P. longipes* (n=4), *P. femoristriga* (n=2), *P. japonicas* (n=2), *P. penicillatus* (n=1), *P. homarus* (n=3), *P. ornatus* (n=4), P. versicolor (n=4), *P. stimpsoni* (n=1), *P. polyphagus* (n=4), and the out-group *T. indicus* (n=2), were used to construct the phylogenetic tree. All members of the family Palinuridae formed distinct clades, separating them from the family Scyllaridae. Among the five species analyzed, *P. polyphagus, P. versicolor*, and *P. ornatus* demonstrated a strong affiliation, with bootstrap support ranging from 80% to 100% within their respective clades. Additionally, *P. longipes* from this study clustered with conspecifics from Sri Lanka and India, receiving 84% support.

In the BLAST search results, *P. longipes* showed 100% query coverage, with similarity percentages of 97% and 98% compared to existing sequences from India and Sri Lanka respectively. The nucleotide base frequencies for *P. longipes* was A: 24.18%, T: 32.86%, C: 20%, and G: 23.06%. The sequence exhibited a strong AT bias, with percentages of 59.38% for *P. longipes*.

The pairwise genetic distance data obtained from four individuals of *P. longipes* showed lowest (0.001) distance between species from Sri Lanka (KF548578) and India (MN863505) and highest (0.212) distance between species from the current study and Vietnam (MN810304) (Table 1).

**Table 1.** Pairwise genetic distance among four individuals of *P. longipes* used to construct the phylogenetic tree. The sequence generated in this study is marked as Bangladesh*.

| Sl No. |  | 1 | 2 | 3 |
| --- | --- | --- | --- | --- |
| 1 | *Panulirus longipes* (L2301SN-01 - Bangladesh*) |  |  |  |
| 2 | *Panulirus longipes* (KF548578 - Sri Lanka) | 0.006 |  |  |
| 3 | *Panulirus longipes* (MN863505 - India) | 0.008 | 0.001 |  |
| 4 | *Panulirus longipes* (MN810304 - Vietnam) | 0.212 | 0.017 | 0.019 |

**Table 2.** The comparative morphological identification features of spiny lobster species collected from Saint Martin’s Island and other four species which were previously reported from Bangladesh.

| Features | *P. longipes* | *P. homarus* | *P. polyphagus* | *P. versicolor* | *P. ornatus* |
| --- | --- | --- | --- | --- | --- |
| Color of carapace | Dark brown with white and orange color spots | Brownish to greenish | Brownish to greenish | Brownish to greenish and black blotches defined with white lines | Greenish blue |
| Cross bands in antennular flagella | Present | Present | Present | Absent | Present |
| Color of frontal horns | Brown with orange tips | Brown with orange tips | Brown with pale white tips | Dark brownish to black with white lines | Dark brownish to black with white lines |
| Spines on antennular plate | 4 separate spines | 4 separate spines | 2 separate spines | 2 pairs of separated unequal spines | 2 pairs of separated spines |
| Spines behind frontal horns | 2 medium spines with 4 conspicuous spines in-between | 3 spines with no spinules in-between | 2 spines with no spinules in-between | 2 medium spines with no spinules in-between | 4 medium spines with no spinules in-between |
| Color of abdominal segments | Dark brown with pale white spots | Olive green | Green with white lines along posterior margin of each somite | Brownish blue with white lines along posterior margin of each somite | Greenish blue with broad transverse dark bands |
| Presence of transverse grooves on abdominal segments | Non-interrupted transverse grooves with a middle notch on 2nd, 3rd and 4th segments | Non-interrupted transverse grooves with crenulated anterior margins | No transverse grooves | No transverse grooves | No transverse grooves |
| Presence of exopod in 2nd maxilliped | Present | Present | Present | Present | Present |
| Presence of exopod in 3rd maxilliped | Present | Absent | Absent | Absent | Absent |

**Table 3.** Updated checklist of lobsters found in Bangladesh. The International Union for Conservation of Nature (IUCN) Red List categories are denoted as EN (Endangered), CR (Critically endangered), VU (Vulnerable), NT (Near threatened), LC (Least concern), DD (Data deficient) and NE (Not evaluated).

| SL no. | Family | Species | English name | IUCN status (National) | IUCN status (Global) | References |
| --- | --- | --- | --- | --- | --- | --- |
| 1 | Palinuridae | *Panulirus versicolor* | Painted spiny lobster | EN | LC | Ahmed *et al.* (2008); Ahmed *et al.* (2022); Habib *et al.* (2023) |
| 2 | Palinuridae | *Panulirus polyphagus* | Mud spiny lobster | VU | LC | Ahmed *et al.* (2008); Ahmed *et al.* (2022); Habib *et al.* (2023) |
| 3 | Palinuridae | *Panulirus longipes* | Long-legged spiny lobster | - | LC | Present study |
| 4 | Palinuridae | *Panulirus homarus* | Scalloped spiny lobster | VU | LC | Ahmed *et al.* (2022); Habib *et al.* (2023) |
| 5 | Palinuridae | *Panulirus ornatus* | Ornate spiny lobster | VU | LC | Ahmed *et al.* (2008); Ahmed *et al.* (2022); Habib *et al.* (2023) |
| 6 | Scyllaridae | *Thenus indicus* | Mud bug | - | DD | Ahmed *et al.* (2023); Habib *et al.* (2023) |
| 7 | Scyllaridae | *Thenus orientalis* | Flathead lobster | NT | LC | Ahmed *et al.* (2008); Habib *et al.* (2023) |

## Discussion

This study was conducted to revise the records of spiny lobster species in Bangladesh by examining the specimen collection from Saint Martin’s Island. A previous study had identified four spiny lobster species from the genus Panulirus in the water of Bangladesh: *Panulirus homarus, P. ornatus, P. polyphagus and P. versicolor*. According to the IUCN Bangladesh (2015), P. versicolor is listed as endangered (EN), while *P. homarus, P. ornatus*, and *P. polyphagus* are assessed as vulnerable (VU) on a national level. However, on a global scale, all of these species are considered to be of least concern (LC) (IUCN, 2021). Globally, *Panulirus longipes* (A. Milne-Edwards, 1868) is classified as a species of least concern (LC), and it has been newly recorded in Bangladesh among the recognized species (IUCN, 2023). Previous studies have utilized both taxonomic and molecular techniques to identify lobsters in Bangladesh (Ahmed et al., 2022; Ahmed et al., 2023). More recently, a new record of the slipper lobster *Thenus indicus* was confirmed (Ahmed et al., 2023). Likewise, new records of spiny lobsters in various regions have been established using morpho-molecular methods (Idreesbabu et al., 2018; Hettiarachchi et al., 2022; Ng et al., 2022; Ng et al., 2023).

The mitochondrial COI genes were used to confirm morphological identifications. Consistent with earlier research, the produced sequences showed a high AT composition throughout the Palinuridae family (Matzen da silva et al., 2011; Jeena et al., 2016; Ahmed et al., 2022). The high rate of G to A mutation (Haag-Liautard et al., 2008) or variations in the selection efficiency across distinct nucleotides could be the cause of this AT bias. Since AT base pairs typically have weaker hydrogen bonds, spontaneous deamination—which can lead to base substitutions—is more likely to occur. As a result, organisms with higher AT content might undergo more mutations, which over time could boost their genetic variety and capacity for adaptation. The minimum genetic distance within the Palinuridae family was observed between *P. homarus* and *P. ornatus* (15.5%), which aligns with the findings of Ahmed et al. (2022) for COI. The maximum genetic distance was recorded between *P. longipes* and *P. ornatus* (25.8%). In contrast, Hettiarachchi et al. (2022) reported the highest genetic distance (41.7%) between *P. homarus* and *P. stimpsoni*, and the lowest (25.8%) between *P. japonicas* and *P. stimpsoni*.

The phylogenetic analysis revealed that intraspecific sequences formed monophyletic groups, confirming the COI gene marker’s effectiveness in species identification and in elucidating phylogenetic relationships among closely related species. All species within the family Palinuridae (represented by five species in our study) clustered into distinct clades, separate from the Scyllaridae family used for comparison. Among these five species, *P. polyphagus, P. versicolor*, and *P. ornatus* showed strong associations with 80-100% bootstrap support within their respective clades. Furthermore, *P. longipes* from the present study grouped with conspecifics from Sri Lanka and India with 84% support. The results also indicated a close relationship between *P. longimanus* and *P. femoristriga*, supporting previous studies (Ptacek et al., 2001; Juinio-Meñez & Ravago et al., 2003). Additionally, Ahmed et al. (2022) had earlier demonstrated the effectiveness of molecular techniques in distinguishing spiny lobsters in Bangladesh.

Identification based on only morphology may lead to misidentification, especially in case of decapod crustaceans. In such cases, DNA barcoding is a great tool for accurate identification of species. This study represents a significant contribution to the understanding of spiny lobster diversity in the Bay of Bengal. Through molecular characterization, we have confirmed the presence of *Panulirus longipes* in Saint Martin’s Island, Bangladesh, a new record for the country. This finding expands the known geographic range of the species and highlights the importance of continued research into the marine biodiversity of this region. Further research is needed to explore the population dynamics, habitat preferences, and ecological role of *P. longipes* in Saint Martin’s Island, Bangladesh. Understanding these aspects will be crucial for effective conservation and management of this valuable species. Additionally, continued molecular studies can help identify other potentially new species or genetic variations within existing populations.

## Author Contribution Statement

Sadia Afrin Kamal: Data curation; methodology; writing- original draft. Nur-A-Zannat: Data curation; methodology; writing- original draft; writing- review and editing. Umme Khadija: writing- review and editing. Kazi Ahsan Habib: Conceptualization; investigation; methodology; supervision; writing- review and editing.

